# Biomechanical Evaluation of Femoral Neck Fracture Fixation Based on Finite Element Analysis

**DOI:** 10.1101/2024.11.30.626173

**Authors:** Laureb Rao, Samarth Mittal, Apoorva Kabra, Vivek Trikha, Anoop Chawla, Kaushik Mukherjee

**Affiliations:** Transportation Research and Injury Prevention Centre, Indian Institute of Technology Delhi, New Delhi, India; Department of Orthopaedics, Jai Prakash Narayan Apex Trauma Centre, All India Institute of Medical Sciences, New Delhi, India; Department of Mechanical Engineering, Indian Institute of Technology Delhi, New Delhi, India

**Keywords:** Finite element analysis, Cannulated Screws, Dynamic Hip Screw (DHS), Femoral neck system, Femoral neck fracture, Pauwels type III fracture, musculoskeletal loading

## Abstract

Pauwels type III fracture, wherein the fracture plane makes an angle of 50°-70° with the transverse plane, is the most unstable femoral neck fracture. Although there are various types of internal fixation devices, including Cannulated cancellous screws (CCS), dynamic hip screw (DHS), dynamic hip screw along with anti-rotation screws (DHS+ARS), and a newly introduced femoral neck system (FNS), patients with Pauwels type III fracture often experience implant failure. There was no study investigating the performance of these internal fixators under physiological musculoskeletal loading. The objective of the present study was to compare the biomechanical performance of CCS, DHS, DHS+ARS and FNS under daily physiological activities like normal walking and stair climbing.

The unstable Pauwels type III fracture was virtually induced in a composite femur bone model and was fixed with all four internal fixators. Second-order tetrahedral elements were used to develop the FE models. The stresses, strains, and deformations of intact and reconstructed femur and implant components were compared to understand the biomechanical efficacy.

FNS showed the highest inferior axial displacement of the femoral head while DHS+ARS exhibited the least inferior axial displacement. Among the four fracture fixation systems, DHS+ARS offered the highest post-fixation stability in terms of fracture gap opening (normal walking: 0.82mm and stair climbing: 0.95mm) and contact sliding, while the performance of single hole locking plated FNS was the worst (normal walking: 2.65mm, stair climbing: 3.72mm). Based on this computational study, DHS+ARS seems to be a better performing system in Pauwels type III fracture fixation.

## 1. Introduction

Incidents of hip fractures are estimated to increase to 2.6M per year by the year 2025 [1,2]. Femoral neck fractures account for nearly 50% of the total incidences of hip fractures occurring across the world [3]. Pauwels type III fracture, wherein the fracture plane makes an angle of 50°- 70° with the transverse plane, is the most unstable fracture [4]. Such fractures typically result in higher shear forces along the fracture plane leading difficulty in achieving post-fixation stability [5]. Consequently, the rate of nonunion is high for such fractures [6].

In younger populations with non-osteoporotic bone, internal fixation is the primary choice for femoral neck fracture treatment [7]. There are various internal fixation devices for treating Pauwels type III fractures. Cannulated cancellous screws (CCS) and dynamic hip screw (DHS) are the two most commonly used internal fixators for treating femoral neck fractures [8,9]. Sometimes, anti-rotation screws (ARS) are also used along with DHS (DHS+ARS) to treat Pauwels type III fractures [10]. Recently, the femoral neck system (FNS) has been introduced for femoral neck fracture fixation [11]. Despite all these advances, patients having Pauwels type III fractures treated with internal fixators often experience complications like nonunion (14%) and osteonecrosis (11%). These complications subsequently lead towards the failure of the fracture reconstruction surgery [12]. Therefore, the choice of the most effective implant for Pauwels Type III fracture fixation remains a matter of contention among the surgeons.

There are many clinical, experimental and finite element (FE) studies on the comparison of various internal fixation devices [6,13–21]. The clinical studies primarily compared the efficacies of these fracture fixation devices in terms of blood loss during the surgery, operating time, Harris Hip Score and other post operative complications such as femoral head necrosis, neck shortening and fracture non-union [6,13,22,23]. The experimental studies, involving cadaveric [14–16] and composite [17,18,24] femoral bones, evaluated post-operative load carrying capacities of fractured femurs treated with various internal fixators like FNS, CCS, DHS etc. There have also been quite a few FE-based computational studies on the success of fractured femur reconstructed with internal fixation devices [25–30]. However, the loading conditions considered in most of these experimental and computational studies were oversimplified. The experimental studies primarily neglected the muscle forces [14,16,31–34] whereas the computational studies [28–30,35–37] did not consider physiological loading corresponding to normal walking and stair climbing. Furthermore, FNS, being a newly introduced implant system, its biomechanical performances yet to be thoroughly investigated with respect to other internal fixators. A recent study [38] compared FNS with CCS, DHS and DHS+ARS considering biomechanical loading conditions. However, this study used shorter unicortical locking screws while bicortical locking screws are used to treat such fractures since bicortical screws offer improved stability than unicortical ones. To the best of authors’ knowledge, there is no study to assess the performance of FNS with respect to CCS, DHS and DHS+ARS for treating Pauwels type III femoral neck fracture under physiological musculoskeletal loading when bicortical locking screws are used.

Hence, the present study is aimed at *in silico* comparative assessment of the following four internal fixation devices for treating Pauwels type III fracture: partially-threaded CCS, DHS, DHS with the anti-rotational screw (DHS+ARS), and single hole locking plated FNS (Figure 1). Bicortical locking screws were considered throughout the study. The specific objective of the study is to compare the stresses, strains and deformations in the reconstructed femur and implant components during daily physiological activities like normal walking and stair climbing.

**Figure 1:**
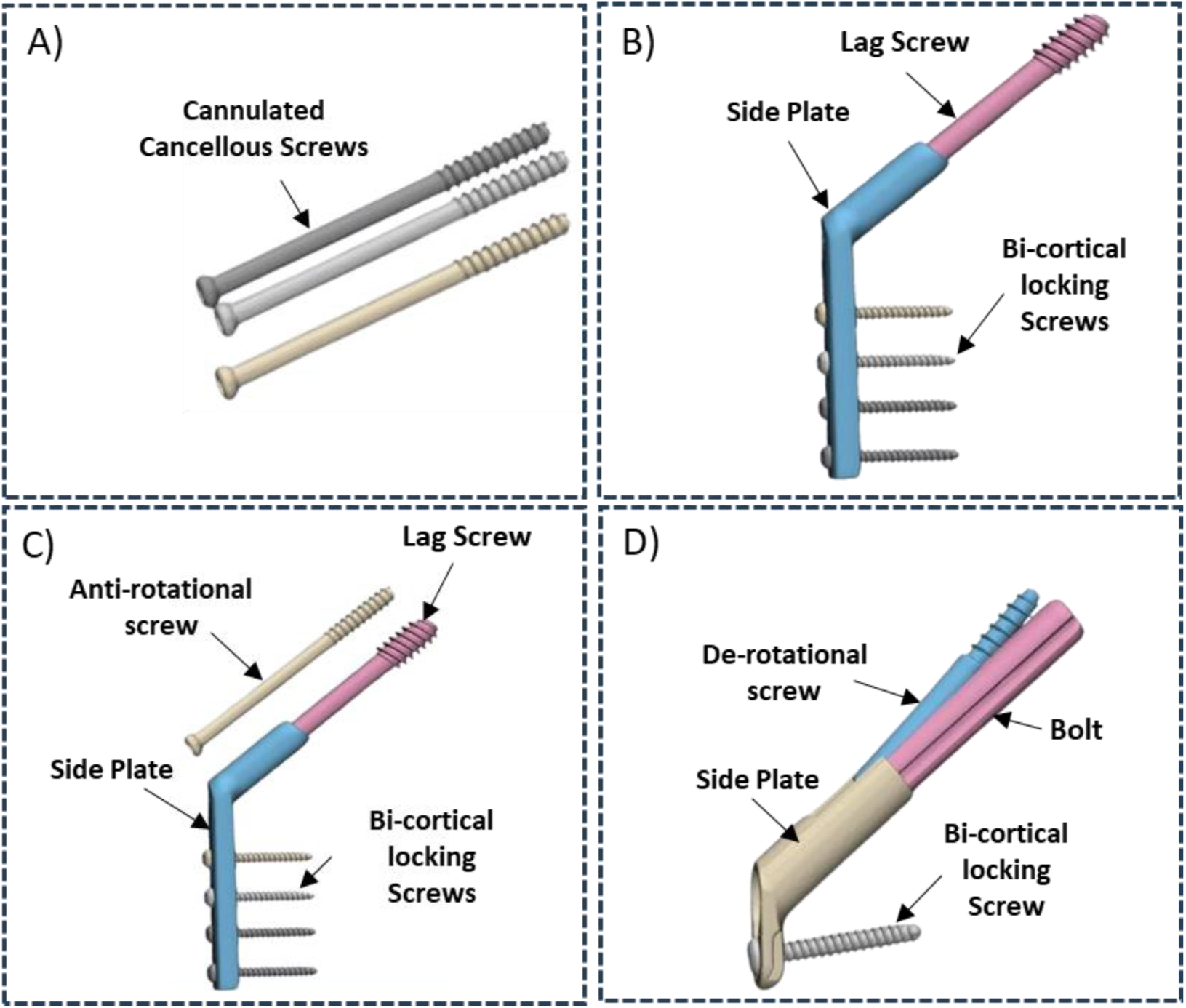
CAD models of the internal fixation devices developed from micro-CT images: A) CCS, B) DHS, C) DHS+ARS and, D) FNS

## 2. Materials and methods

### 2.1 Intact femur model generation and validation

The CAD model of manufacturer-provided fourth-generation composite left femoral bone (Model #3406 Saw Bones, A Pacific research laboratory, Vashon, WA, USA) was used in this study (Figure 2A) to avoid patient-specific variation in geometry and material properties of the femur [39]. The FE model of intact femur was developed from the CAD model in Altair Hypermesh 2020.1 (Altair Engineering Inc., Troy, Michigan, USA) using ten-noded tetrahedral elements. The composite femur model had distinct cancellous and cortical layers which shared common interfacial nodes. Both these layers were considered homogeneous, linear elastic and isotropic [40,41].

**Figure 2:**
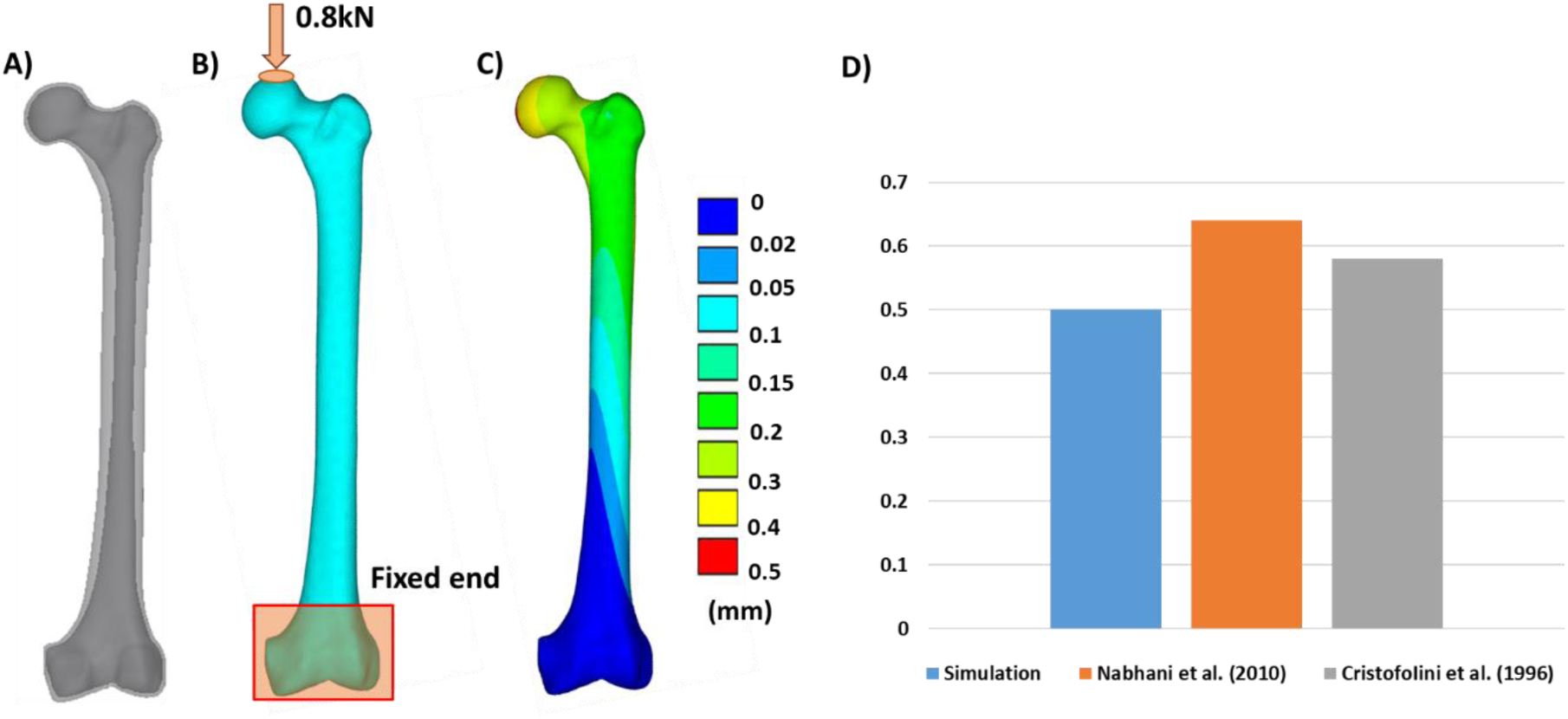
Validation of FE model generation procedure: A)CAD model of intact bone, B) FE model of intact bone with loading and boundary conditions, C) vertical displacement of the femur under prescribed loading, D) comparison of inferior axial displacement of femur with previous studies

The mesh convergence study was conducted on the intact femur by generating meshes with three different element sizes of 2mm, 3mm and 5mm, resulting in 175713, 91258 and 13902 elements, respectively. The difference in the peak von-Mises stress between the coarse and the mid-size mesh was approximately 5%, and that between the mid and the fine size mesh was close to 1%. An element size of 3mm was, therefore, chosen for further analysis as additional mesh refinement did not significantly improve results.

To validate the FE model generation procedure, the FE predicted results were compared with previously published experimental studies [42,43]. For this validation, the FE model of the intact femur was oriented at an abduction angle of 11^0^. A downward compressive force of 800N corresponding to an average body weight of 80kg at mid stance of the gait cycle was applied at the femoral head (Figure 2B). Similar to earlier studies [42,43], 0.5 – 0.6 mm inferior axial displacement of the femoral head was observed (Figure 2C). These qualitative comparisons provided confidence in our FE modelling approach.

### 2.2 Implant-bone assembly modeling

#### 2.2.1 CAD model generation of implant-bone assembly

Pauwels type III fracture was induced virtually in the composite femur bone (Model #3406 Saw Bones, A Pacific research laboratory, Vashon, WA, USA) wherein the fracture plane was at an angle of 70° with respect to the transverse plane (Figure 3A). The CAD models of all the fracture fixation devices were developed from Depuy Synthes (Johnson & Johnson, Raynham, MA, USA) implants. The implants were first scanned (slice thickness: 60 μm, pixel size: 60μm; width*height: 553*629) using micro-CT scanner (Scanco Xtreme CT-II, Scanco Medical AG, Brüttisellen, Switzerland) (Figure 1, 3B). The micro-CT Images were, thereafter, converted into 3D models using Mimics^TM^ V21.0 (Materialise, Belgium). The fractured bone model was assembled with each of these implants (CCS, DHS, DHS+ARS, FNS) to simulate anatomical reduction and internal fixation (Figure 3C). Inputs from the clinicians were considered for placing these implants during this virtual fracture fixation surgery. The tip apex distance (TAD) for all the implants was assumed to be 20mm. On the coronal plane, the first CCS was placed right above the calcar region. The second CCS was placed in the supero-posterior direction with respect to the first one, whereas the third CCS was placed in the supero-anterior direction with respect to the first [44]. The DHS and single hole locking plated FNS were placed in the middle of the femoral head when viewed in the coronal and sagittal planes. All the fracture fixation devices were placed parallel to the femoral neck axis (Figure 3C) following standard surgical protocol. Due to the difference between the CCD angle (caput-collum-diaphysis i.e. femoral neck shaft) of FNS (130 Degree) and sawbone femur (115 Degree), orienting the bolt axis along the femoral neck axis caused a gap of ∼4.5mm between the distal tip of FNS side plate and femoral shaft (Figure 3C).

**Figure 3:**
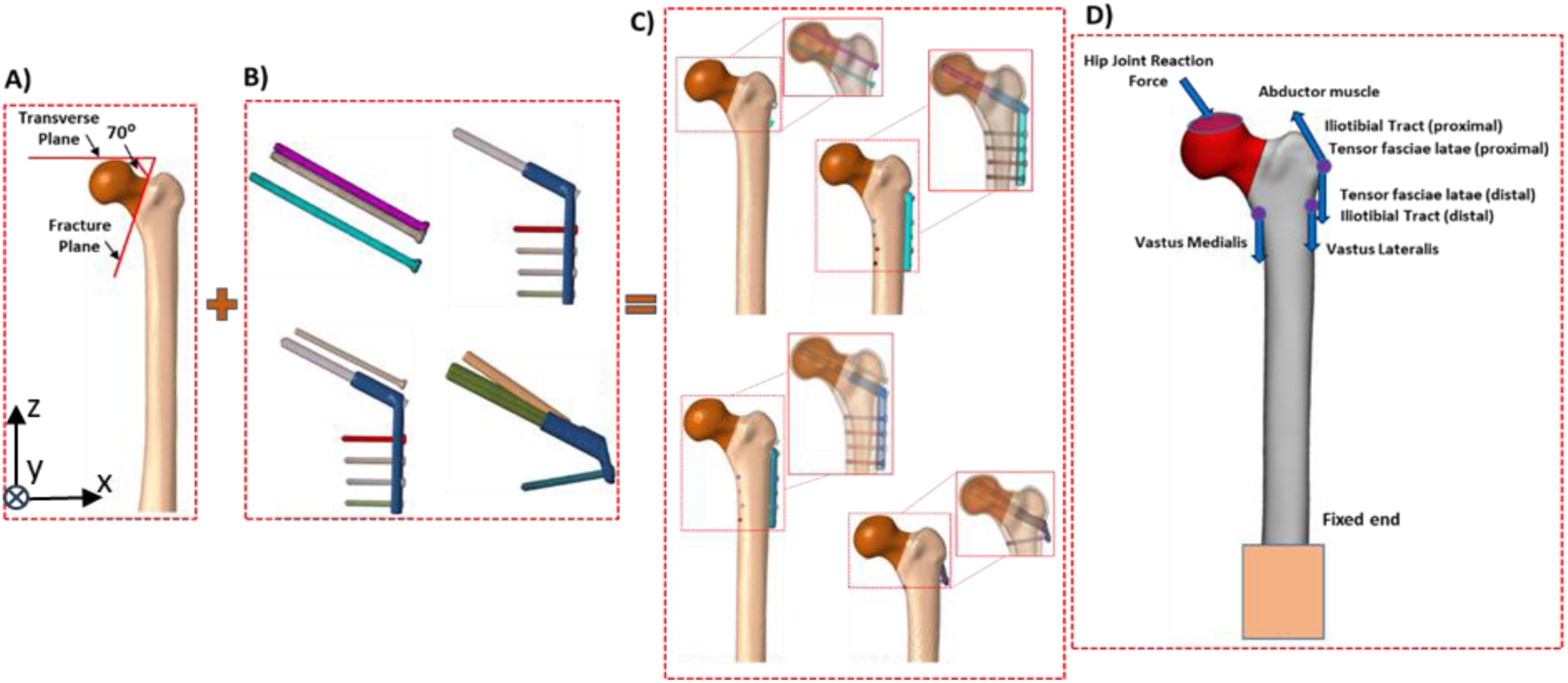
Development of implant-bone FE model: A) femur with Pauwels type III fracture, B) FE models of implants, C) Post-fixation implant-bone FE model D) Forces and BCs for FE analysis used for the simulation of two day to day activities of normal walking and stair climbing

#### 2.2.2 FE model generation

The STL files of the implants and bones were meshed in Altair Hypermesh 2020.1 (Altair Engineering Inc., Troy, Michigan, USA) using ten-noded tetrahedral elements (Figure 3). The threaded portion of the screws was assumed to be cylindrical to reduce computational complexity (Figure 3B). Accordingly, four FE models with CCS, DHS, DHS+ARS and FNS implants were generated for the fractured bone. While for femurs, an element size of 3mm was used, for the implants an element size of 1mm was used for discretization. The resultant number of elements and nodes in all the models are mentioned in Table 1.

**Table 1.** The total number of elements and nodes in intact, CCS, DHS, DHS+ARS and FNS fracture fixation models.

|  | Intact | CCS | DHS | DHS+ARS | FNS |
| --- | --- | --- | --- | --- | --- |
| Number of elements | 67123 | 225286 | 462755 | 504909 | 368407 |
| Number of nodes | 98164 | 331519 | 663563 | 749728 | 560274 |

A frictional interface (coefficient of friction, µ = 0.46) was assumed between the two femoral segments [45]. The interface between the bone and the plate was modelled as frictional contact with µ=0.3 [46]. A bonded interfacial condition was assumed between the threaded portion of the implant systems and bone, whereas a frictional interfacial condition (µ=0.3) was considered between the non-threaded portion of the implants and bone [47]. Cortical locking screw heads of DHS and FNS, being threaded, were modeled as bonded with the side plates. Frictional sliding contact with no separation (with µ=0.3) was assumed between the lag screw and side plate of DHS and the bolt and side plate of FNS [48]. The threaded interface between the de-rotational screw and the bolt of FNS was modelled as bonded. The implants were assumed to be made of Ti6Al4V [41]. All the components were modelled as homogeneous, linear elastic and isotropic (Table 2).

**Table 2.** Material properties for Cortical, Cancellous Bones and the Implants.

|  | Young's Modulus (MPa) | Poisson's ratio |
| --- | --- | --- |
| Cortical Bone [45] [49] | 16800 | 0.3 |
| Cancellous Bone [45][49] | 155 | 0.2 |
| Implants (Ti<sub>6</sub>Al<sub>4</sub>V)[45] | 105000 | 0.3 |

The physiological musculoskeletal loading conditions (Table 3), representing normal walking and stair climbing, were considered here [50,51]. The muscle forces were distributed over a patch of 40-50 nodes representing each muscle attachment site (Figure 3D). The joint reaction force was distributed over a patch of 130 nodes on the femoral head such that the resultant force passed through its center. All the musculoskeletal forces were calculated corresponding to a person with 80kg of body weight [52].

**Table 3.** Various muscle forces acting in normal walking and the stair-climbing conditions [52,53].

| Loading condition | Location of force (muscle) | $F_x$ (N) | $F_y$ (N) | $F_z$ (N) | $F_{Resultant}$ (N) |
| --- | --- | --- | --- | --- | --- |
| Normal Walking | Hip Joint | 423.79 | 257.41 | -1798.78 | 1865.78 |
|  | Abductor muscle | -455.184 | -33.74 | 678.85 | 817.99 |
|  | Tensor fasciae latae (proximal) | -56.51 | -91.03 | 103.59 | 149.27 |
|  | Tensor fasciae latae (distal) | 3.92 | 5.49 | -149.11 | 149.27 |
|  | Vastus Lateralis | 7.06 | -145.18 | -729.08 | 743.44 |
| Stair Climbing | Hip Joint | 465.38 | 475.58 | -1854.48 | 1970.24 |
|  | Abductor muscle | -550.14 | -226.02 | 666.29 | 893.10 |
|  | Iliotibial Tract (proximal) | -82.40 | -23.54 | 100.45 | 132.01 |
|  | Iliotibial Tract (distal) | 3.924 | 6.27 | -131.84 | 132.08 |
|  | Tensor fasciae latae (Proximal) | -24.32 | -38.45 | 22.75 | 50.85 |
|  | Tensor fasciae latae (distal) | 1.57 | 2.35 | -51.01 | 51.09 |
|  | Vastus Lateralis | 17.26 | -175.79 | -1060.26 | 1074.86 |
|  | Vastus Medialis | 69.06 | -310.78 | -2096.2 | 2120.21 |

### 2.3 Criteria for Biomechanical evaluation

Implant material (Titanium alloy) being ductile in nature, von-Mises stresses in implants were considered to assess its failure risks [30]. For bone, the maximum principal strain was considered as the failure criterion since earlier cadaveric and computational studies showed that maximum principal strains could successfully predict the risk of failure [54]. Post-operative stability of implant-bone construct and the micro-movements at the fracture surface govern the fracture healing and thus, influence the success of the surgery [55,56]. Consequently, the following parameters were considered for comparing the biomechanical performance of the four fracture fixation methods:

i. Total displacement of femur and inferior axial displacement of the most superior point of the femoral head,
ii. gap opening and contact sliding of the fracture surfaces (Figure 4)

**Figure 4:**
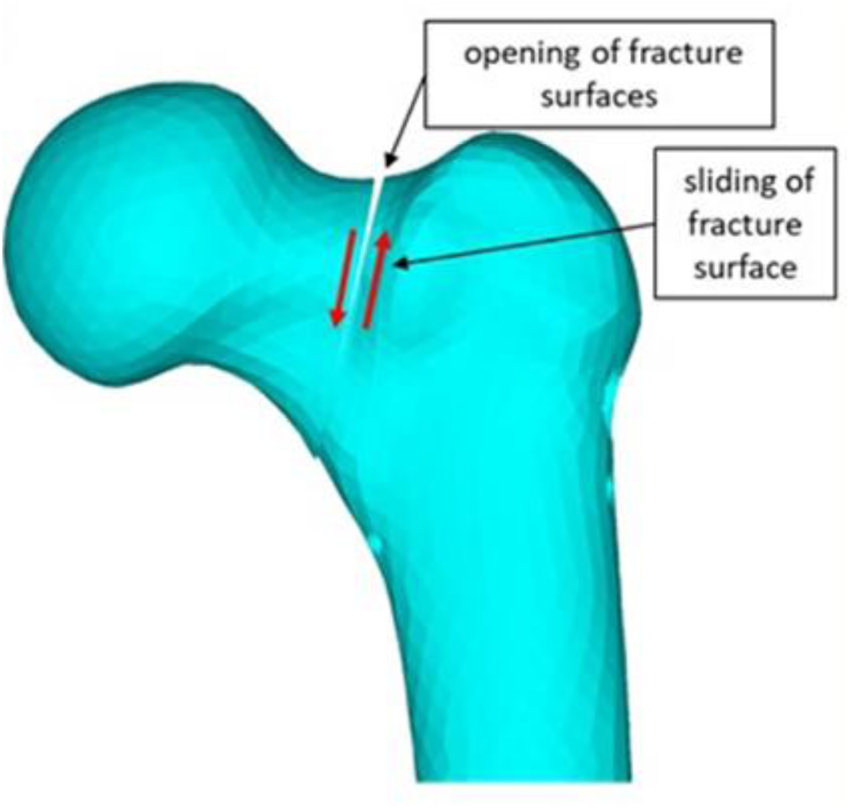
The illustration of opening of fracture surface and the sliding of the fracture surface

## 3. Results

ANSYS Mechanical Product Launcher (APDL) was used to solve the FE models, and the contour plots were obtained to analyze the results. The results based on criterias used for the biomechanical performance evaluation of different implant systems are described below.

### 3.1 Maximum principal strains in femur

The contour plots of the maximum principal strains are shown in Figure 5. For both normal walking and stair climbing cycles, there were differences in strain distributions in proximal region between the intact and implanted femurs. However, maximum principal strains in femur were always less than the failure limits of cortical and cancellous bones for all the models under both loading conditions. The superior calcar region shows reduced principal strains (80%-85%) in all the implanted bone as compared to the intact femur. However, the strains were higher in the inferior calcar region (30%-40%). The greater trochanteric region of the implanted bone experienced less strains compared to the intact bone (20%-30%).

**Figure 5:**
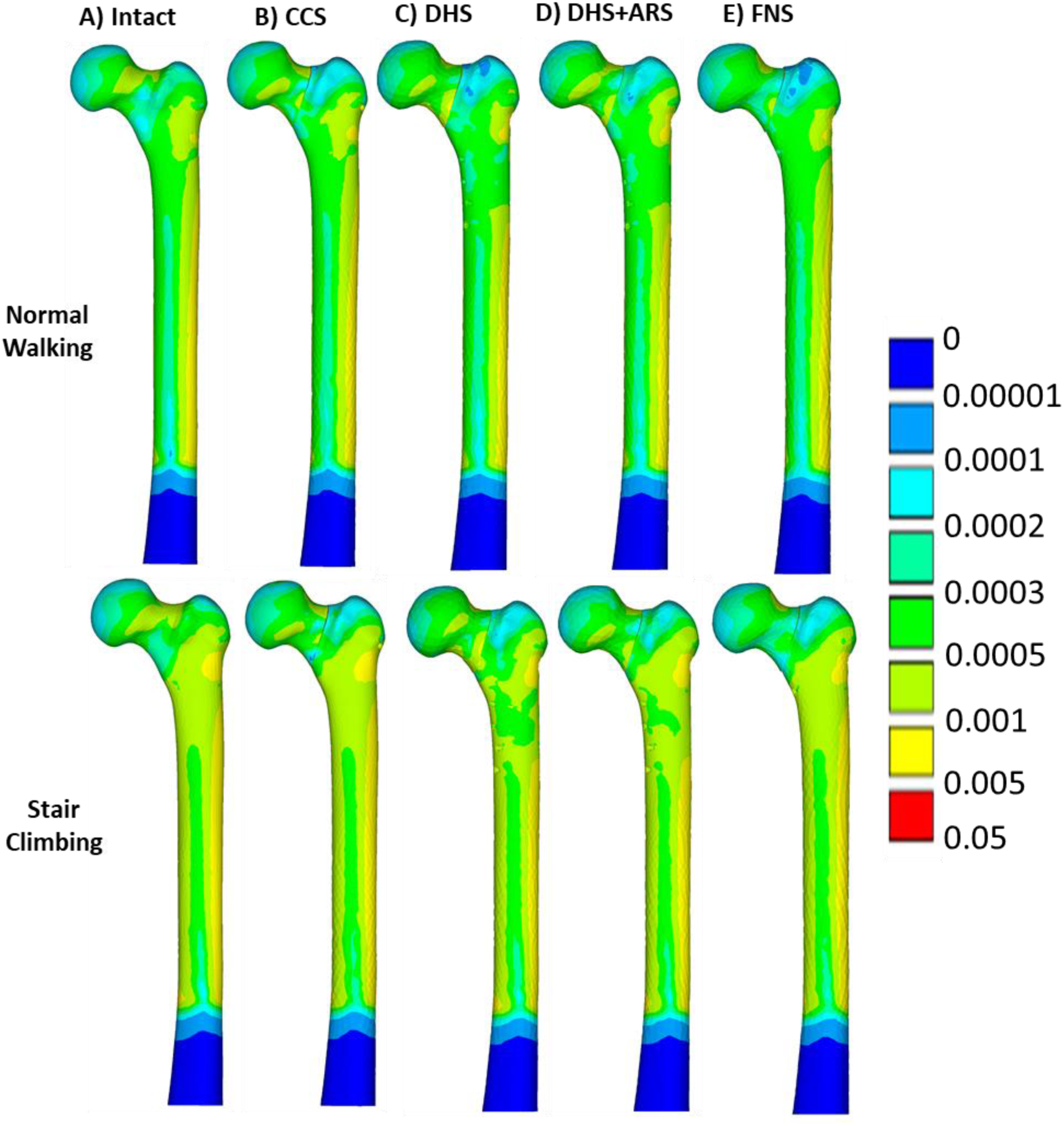
Maximum principal strain distribution in the bone subjected to the loading conditions of normal walking and stair climbing in the case of: A) Intact bone, B) CCS fixation, C) DHS Fixation, D) DHS+ARS Fixation

### 3.2 Von Mises stresses in implant

Under both loading conditions, contour plots for von-Mises stress were plotted for the intact femur and fractured femur fixed with CCS, DHS, DHS+ARS and single hole FNS (Figure 6). The maximum values of the Von-Mises stress for all the FE models are given in Table 4. The maximum von-Mises stress in CCS occurred near the fracture plane (Figure 6). In DHS, the maximum von-Mises stress was observed in the lag screw near the fracture plane (Figure 6). In FNS fixation, the maximum von-Mises stress was present at the junction of the locking screw and the side plate (Figure 6). Furthermore, for DHS and FNS systems, the von-Mises stress in lag screw and bolt, respectively, was significantly higher near the fracture plane. For all the models, stresses in stair climbing condition were higher than normal walking condition (Table 4, Figure 6). It should, however, be noted that these von Mises stresses never exceeded the yield point of the Titanium alloy (Ti-6Al-4V).

**Figure 6:**
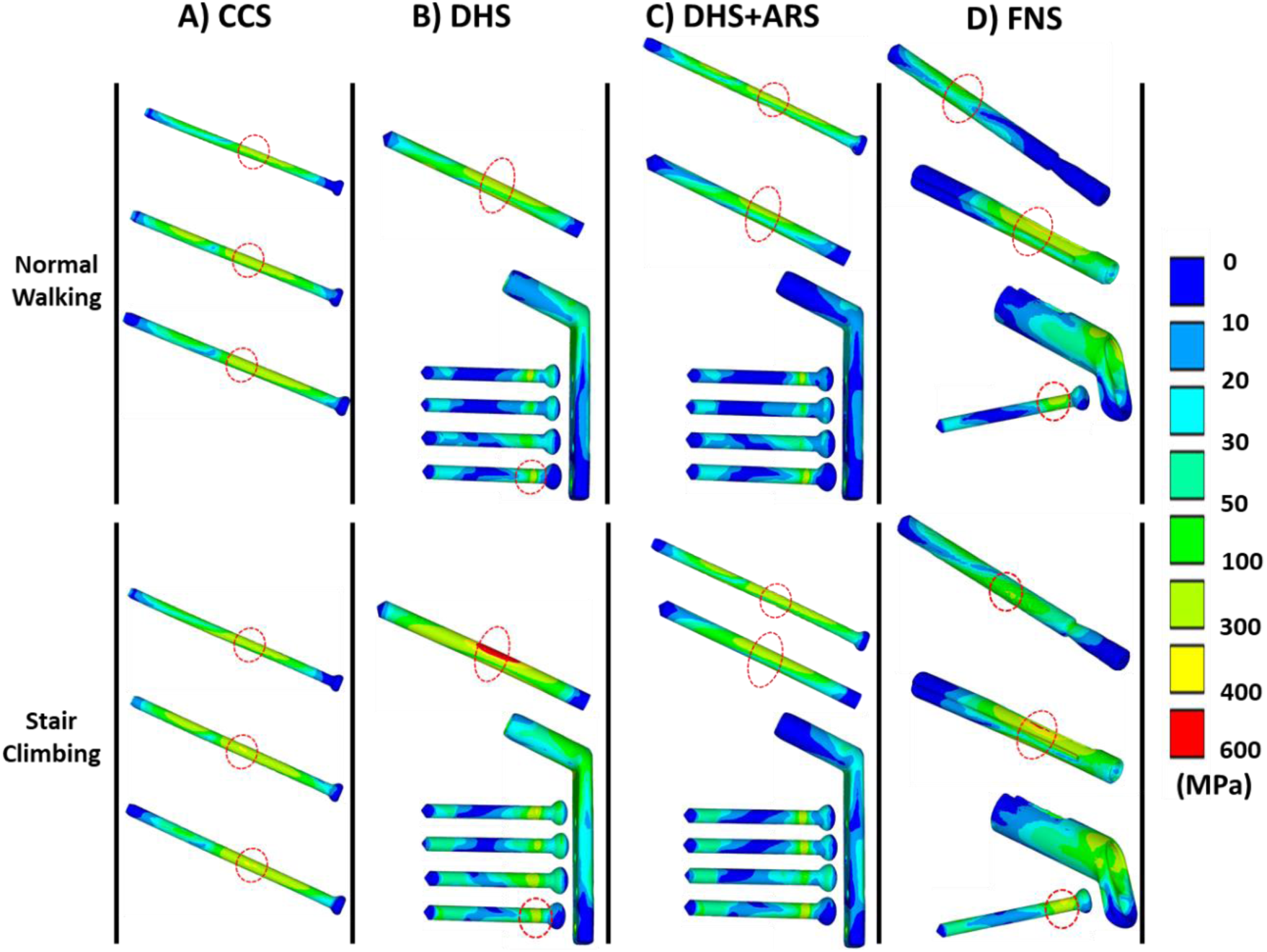
von-Mises stress distribution in the implants subjected to the loading conditions of normal walking and stair climbing in the case of: A) CCS fixation, B) DHS Fixation, C) DHS+ARS Fixation, and D) FNS, (The encircled region shows the location of maximum)

**Table 4.**
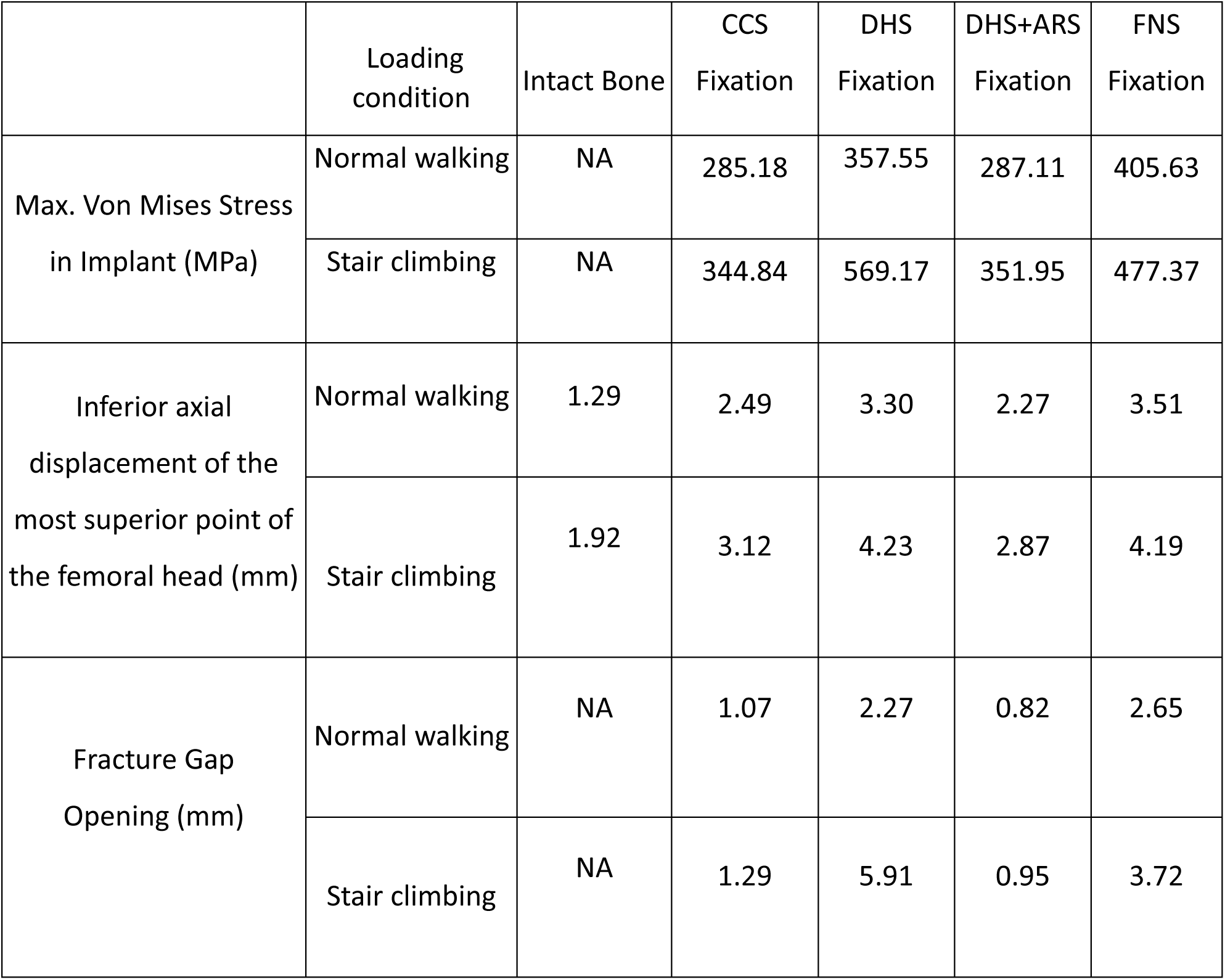
Maximum values of Von-Mises Stress in bone, Von-Mises Stress in implant, vertical displacement of femoral head and the fracture gap opening.

|  | Loading condition | Intact Bone | CCS Fixation | DHS Fixation | DHS+ARS Fixation | FNS Fixation |
| --- | --- | --- | --- | --- | --- | --- |
| Max. Von Mises Stress in Implant (MPa) | Normal walking | NA | 285.18 | 357.55 | 287.11 | 405.63 |
|  | Stair climbing | NA | 344.84 | 569.17 | 351.95 | 477.37 |
| Inferior axial displacement of the most superior point of the femoral head (mm) | Normal walking | 1.29 | 2.49 | 3.30 | 2.27 | 3.51 |
|  | Stair climbing | 1.92 | 3.12 | 4.23 | 2.87 | 4.19 |
| Fracture Gap Opening (mm) | Normal walking | NA | 1.07 | 2.27 | 0.82 | 2.65 |
|  | Stair climbing | NA | 1.29 | 5.91 | 0.95 | 3.72 |

### 3.3 Displacement of femoral head

Out of all the four internal fixators, FNS exhibited the highest inferior axial displacement of the most superior point of the femoral head, whereas DHS+ARS resulted in the lowest displacement (Table 4) for both the loading conditions. A similar trend was evident from contour plots of total displacements of intact and implanted femurs (Figure 7). It should, however, be noted that the deformation in the intact femur was less than all the fractured femurs with internal fixators (Table 4, Figure 7). Furthermore, the displacement of the femoral head was higher in stair climbing as compared to normal walking.

**Figure 7:**
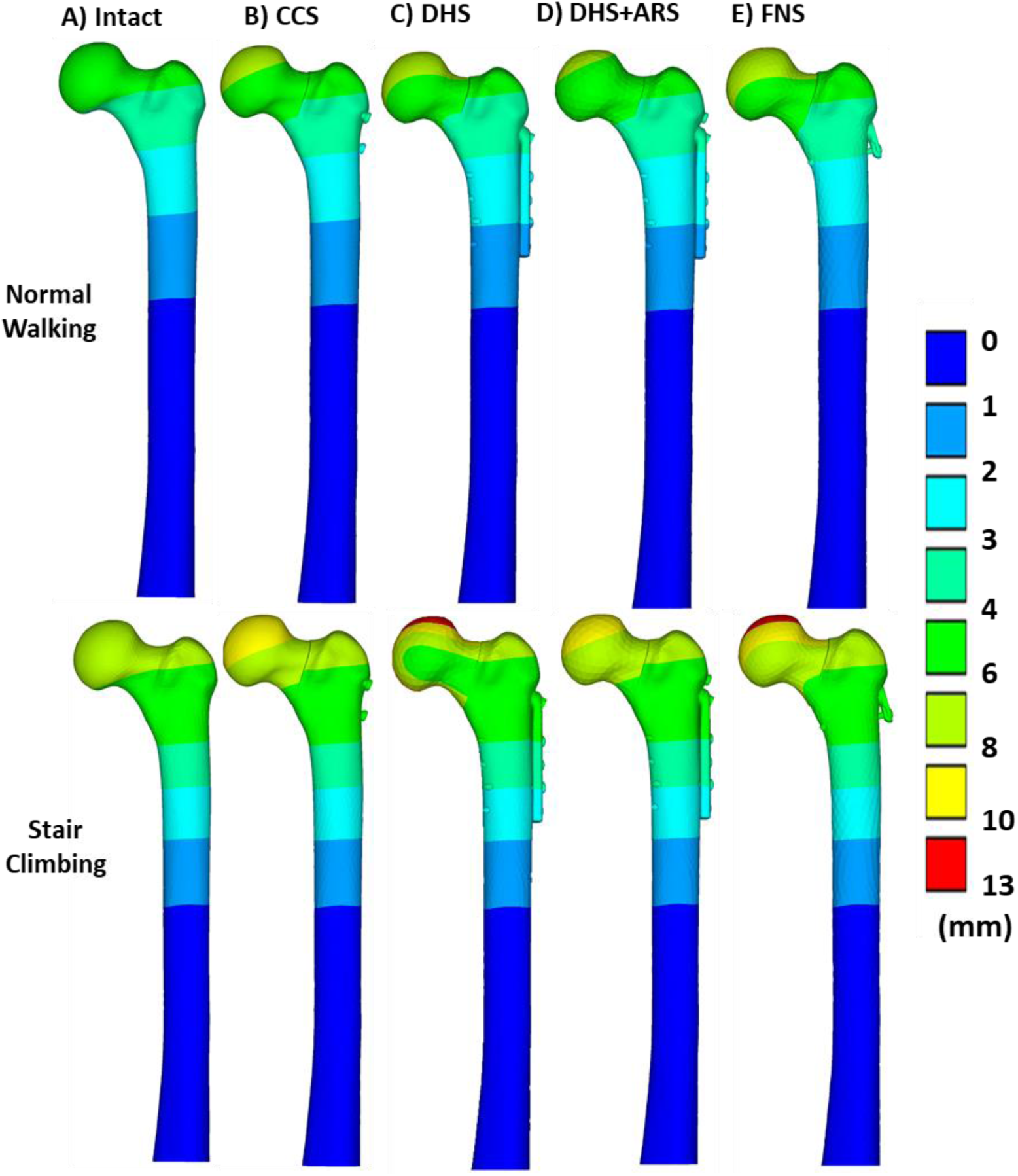
Total displacement of the femoral head in the implant bone assembly subjected to the loading conditions of normal walking and stair climbing in the case of: A) Intact bone, B) CCS fixation, C) DHS Fixation, D) DHS+ARS Fixation, and E) FNS

### 3.4 Gap opening and contact sliding of fractured surface

For all four fractured models, maximum gap opening was observed at the superior calcar region of the fractured femur. It was observed that CCS fixation resulted in minimum gap opening while DHS+ARS exhibited minimum contact sliding (Table 4, Figure 8). DHS fixation, on the contrary, caused maximum gap opening as well as contact sliding under both loading conditions. Both gap opening and contact sliding were found to be higher during stair climbing than that of normal walking condition.

**Figure 8:**
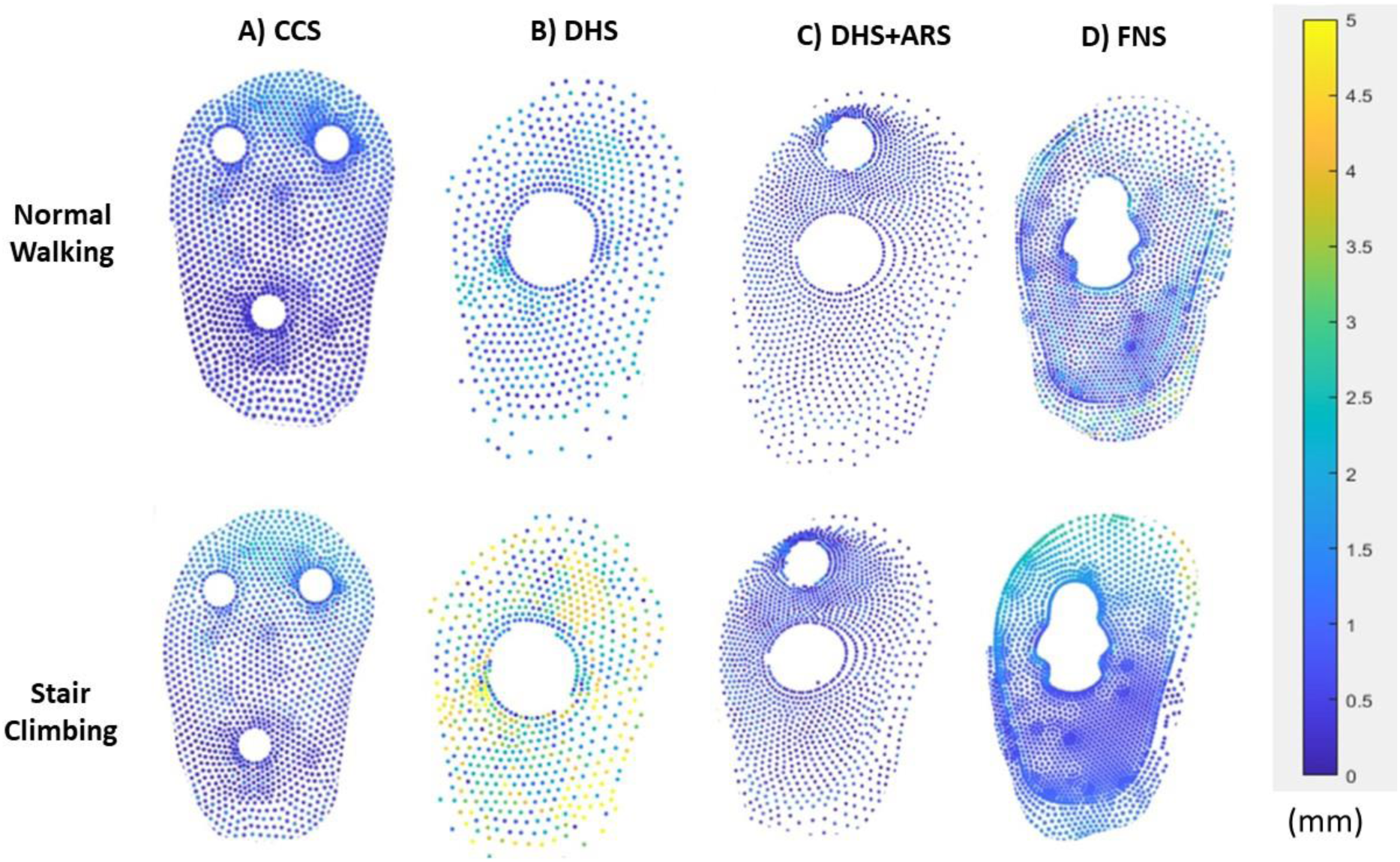
Contact sliding between the two fracture fragments under the normal walking and stair climbing loading conditions in case of A) CCS fixation, B) DHS fixation, C) DHS+ARS fixation, and D) FNS

## 4. Discussions

Earlier studies showed that the Pauwels type III fractures have the highest rate of operative complications [57]. Higher shear forces in the case of Pauwels type III fracture make it difficult to manage this type of fracture [58]. Earlier studies have also evaluated the biomechanical performance of different internal fixation techniques. Zhirong et al. (2021) [29] performed a study to compare the FNS with three CCS arranged in different configurations. Their findings indicated that the three CCS provided greater biomechanical stability than the FNS. Jun-ki Moon et al. (2022)[33] conducted an experimental study comparing FNS and DHS to conclude that both FNS and DHS showed comparable performance. The present study was aimed at the biomechanical evaluation of four fracture fixators, namely, CCS, DHS, DHS+ARS and FNS, for treating Pauwels type III femoral neck fracture. *In silico* models were developed for each reconstruction, and physiological loading of normal walking and stair climbing conditions were considered for further analysis. Stresses, strains and deformations in reconstructed femurs and implant components were compared for biomechanical assessment.

Based on the FE results obtained it can be concluded that overall, there was a reduction in maximum principal strains in the superior calcar region and greater trochanteric region of all the implanted bones as compared to the intact femur. However, the inferior calcar region of the implanted bone experienced higher strains as compared to the intact femur. The reduced principal strain distribution near the neck and the greater trochanteric region indicates stress shielding in the implanted femur. Stress shielding in the femoral neck region was also documented in the previous studies [59,60], and therefore We L. et al. (2022) [60] suggested the removal of the implants after the fracture healing. Ning Li et al. (2024) [61] also indicated that the inferior calcar region of the femur, experienced higher strains in the commonly used implants such as CCS and DHS. The von-Mises stresses in all four implant components were less than yield limit of the Titanium alloy (Ti6Al4V), i.e. 950 MPa [62]. However, DHS implant component always experienced the highest von Mises stresses, whereas DHS+ARS implant components exhibited the least von Mises stresses under physiological conditions. It should, however, be noted that for all four implant systems, higher stresses were observed near the fracture site (encircled in Figure 6). The higher stress distribution near the fracture site in the CCS and DHS fixation was well corroborated with the previous studies [63,64].

In femoral neck fractures, the primary function of the implants is to stabilize the femoral head and restore normal anatomy of the femoral neck to achieve optimal stability. The stability of the femoral head is also critical because any displacement or misalignment of the femoral head can lead to significant complications such as avascular necrosis, non-union or malunion of the fracture [65,66]. Therefore, the best system of fracture fixation should minimize total displacements of implant bone assembly under physiological loading condition. Hence, to assess the stability of the femoral head, the following four parameters were considered for comparison: total displacement and inferior axial displacement of the most superior point of the femoral head, gap opening and contact sliding of the fractured surfaces. The fracture gap opening for the four fixation techniques indicated that the femoral head experienced a rotation about the inferior calcar region under physiological loading. Among the four fracture fixation techniques considered in this study, DHS+ARS provided the best results in terms of displacements of the femoral head, fracture gap opening and contact sliding under both normal walking and stair climbing loading conditions. In comparison, single-hole locking plated FNS performed the worst for all these parameters under both physiological loading conditions.

The overall findings of the current study are consistent with the earlier studies. Previous computational studies [67,68] showed that in the case of Pauwels type III fracture having a fracture angle of 70°, FNS exhibited higher total displacement as compared to CCS system under simplified loading regime. Relatively higher stresses around the fracture plane were also reported in these studies [67,68]. Furthermore, multiple clinical [13,69] and computational studies [70] have established the biomechanical advantage of using DHS or DHS+ARS over CCS for treating Pauwels type III fractures. Several earlier studies have demonstrated the inferiority of FNS over CCS, DHS and DHS+ARS which also corroborates well with the findings of the current study [35,71–74]. Although a simplified loading regime was considered in previous computational studies, while the present study considered physiological musculoskeletal loading conditions, the results of the present study exhibited similar trends. These corroborations with previous studies provided further confidence in the inferences drawn in this study.

There are certain limitations of the present finite element study. In this FE study, the detailed thread modelling was neglected since the focus was on understanding the mechanical behaviour of the femoral reconstruct under physiological loading. Hence, the influence of screw thread geometry on the overall stability was out of the purview of this study. Although the results of the present study were qualitatively validated against earlier studies by [37,68,70,75], direct one-to-one validation of FE predicted results were out of the scope of this present study. Furthermore, the simulated Pauwels type III fracture was assumed to be a simple transcervical fracture, neglecting any irregularity of the fracture plane. Other clinical parameters such as the operating time, blood loss during the surgery, and implant footprint on the bone were out of the scope of this study.

It should be noted that the thread length of de-rotational screw of femoral neck system is smaller than that of CCS and ARS. Thread lengths are known to influence implant stability [36,76]. Additionally, the gap between the FNS side plate and femoral bone might also compromise the biomechanical performance of FNS. Further studies are envisaged to investigate these factors thoroughly. Limitations notwithstanding, this study provides a computational platform to compare the biomechanical efficacies of four fracture fixation implants (CCS, DHS, DHS+ARS and FNS) under physiological loading. Furthermore, this study also provides key insight regarding regions with high stresses for all these implant components under physiological loading conditions.

## 5. Conclusions

This biomechanical study compared four fracture fixation implants (CCS, DHS, DHS+ARS and FNS) for treating Pauwels type III fracture with a fracture angle of 70°. Musculoskeletal forces corresponding to two physiological activities, i.e. normal walking and stair climbing, were considered. Among the four fracture fixation systems, DHS+ARS offered the highest post-fixation stability while the performance of single hole locking plated FNS was the worst. In all four cases, higher stresses were observed near the fracture plane. However, stresses in none of the four fixation systems exceeded the yield limit of Ti6Al4V. Hence, based on this computational study, DHS+ARS seems to be a better performing system for treating Pauwels type III fracture with a fracture angle of 70°.

